# A common pattern of DNase-I footprinting throughout the human mtDNA unveils clues for a chromatin-like organization

**DOI:** 10.1101/193037

**Authors:** Amit Blumberg, Charles G. Danko, Anshul Kundaje, Dan Mishmar

## Abstract

Human mitochondrial DNA (mtDNA) is believed to lack chromatin and histones. Instead, it is coated solely by the transcription factor TFAM, which binds the mtDNA without sequence specificity and packs it into a bacterial-like nucleoid in a dose-dependent fashion. We asked whether mtDNA packaging is more regulated than once thought. As a first step to address this question, we analyzed mtDNA DNase-I-seq experiments in 324 different human cell types and found, for the first time, a pattern of 29 Genomic footprinting (DGF) sites throughout the mtDNA shared by ∼90% of the tested samples. Low SNP density at the DGF sites, and their conservation in mouse DNase-seq experiments, reflect strong selective constraints. Co-localization of the DGFs with known mtDNA regulatory elements and with recently-discovered transcription pausing sites, suggest a role for such DGFs in mtDNA transcription. Altered mtDNA DGF pattern in IL-3 treated CD+34 cells offer first clue to their physiological importance. Taken together, human mtDNA has a conserved and regulated protein-DNA organization, which is likely involved in regulation of mtDNA gene expression.

## Introduction

Global regulation of gene expression in the human genome is governed by a combination of chromatin structure, its availability to the transcription machinery, histone modifications and DNA methylation status (Roadmap Epigenomics et al.2015; Zhu and Guohong 2016). However, this general scheme of gene expression regulation does not apply to the only component of the human genome that resides in the cytoplasm – the mitochondrial genome (Bogenhagen 2012).

The mitochondrion has a pivotal role in cellular ATP production via the oxidative phosphorylation system (OXPHOS). Because of its centrality to life, OXPHOS dysfunction leads to devastating diseases, and plays a major role in common multifactorial disorders (Marom et al. 2017). In the vast majority of eukaryotes, OXPHOS protein-coding genes are divided between the mitochondrial and nuclear genomes (mtDNA and nDNA, respectively) with most (∼80) in the latter (nDNA), and 13 in the former (Calvo and Mootha 2010). Unlike the nDNA, mtDNA gene expression is currently thought to be regulated by a relatively simple system with relic characteristics of the ancient bacterial ancestor of the mitochondria (Gustafsson et al. 2016). Accordingly, the core regulators of mtDNA transcription form an evolutionarily conserved set of factors including two transcription factors (TFAM, mtTF2B) (Falkenberg et al. 2002; Cotney et al. 2007; Shutt et al. 2010; Shi et al.2012), one RNA polymerase (POLRMT) (Gaspari et al. 2004), one termination factor (MTERF1) (Yakubovskaya et al. 2010; Guja and Garcia-Diaz 2012), and a single known elongation factor TEFM (Minczuk et al. 2011). Secondly, some core mtDNA transcription regulators have bacterial characteristics, such as the phage ancestral structure of POLRMT (Ringel et al. 2011). Thirdly, mtDNA genes are co-transcribed in strand-specific polycistrons (Aloni and Attardi 1971). Specifically, 12 mRNAs encoding protein subunits of the oxidative phosphorylation system (OXPHOS), 14 tRNAs and 2 ribosomal RNA genes are encoded by the heavy strand; the mtDNA light strand encodes for a single mRNA (the ND6 subunit of OXPHOS complex I) and 8 tRNA molecules. The polycistronic transcripts of both strands are, in turn, cleaved into individual (or di-cistron) transcripts following the tRNA punctuation model (Ojala et al. 1981; Montoya et al. 2006). Finally, it has been assumed that regulatory elements of human mtDNA gene expression are mainly located within the major mtDNA non-coding region, the D-loop. Nevertheless, growing body of evidence suggest that mtDNA transcriptional regulation is more complex than once thought. Firstly, it has been found that known regulators of nuclear genes transcription, such as MEF2D (She et al. 2011), glucocorticoid receptor (Demonacos et al. 1993), the mitochondrial receptor for the thyroid hormone tri-iodothyronine (Wrutniak et al. 1995) bind the human mtDNA and regulate its gene expression (Leigh-Brown et al. 2010; Szczepanek et al. 2012). Secondly, the usage of genomics tools, such as chromatin immunoprecipitation (ChIP)-seq, enabled us and others the identification of nuclear transcription regulators (C-Jun, JunD and CEBPb) that in vivo bind the human mtDNA (Blumberg et al. 2014; Marinov et al. 2014) and imported into the mitochondria (Blumberg et al. 2014). As the binding sites of such factors occur within the mtDNA coding region, it is possible that human mtDNA coding sequences are written in two languages – the gene-coding one and the regulatory one. These pieces of evidence led us to hypothesize that the two billion years of endosymbiosis, and subsequent multiple adaptation events of the OXPHOS to changing energy requirements (Ellison and Burton 2010; Bar-Yaacov et al. 2015; Scott et al. 2015), were also accompanied by adaptation of mtDNA gene expression regulation to the eukaryote host environment.

If mtDNA regulation has converged, at least in part, into the gene regulatory system of the host, is it possible that the mtDNA regulatory system developed chromatin-like organization? In contrast to histone coating, and chromatin structure that modulates nuclear gene expression, the mtDNA higher order organization, the nucleoid, is considered far less complex (Brown et al. 2011). Specifically, the mtDNA is known to be coated by a single HMG box protein, the transcription factor TFAM, which binds the mtDNA without any sequence specificity (Kanki et al. 2004; Kaufman et al. 2007; Kukat et al. 2015). Recently, a study employing a combination of high resolution microscopy and cell biology techniques revealed that TFAM coats the mtDNA in a dose-dependent manner, and that TFAM molecules bind the mtDNA every ∼8bp (Kukat et al. 2015). This observation is consistent with recent analysis of TFAM ChIP-seq experiments in HeLa cells (Wang et al. 2013). All these findings support a very simple higher order organization of the mammalian mtDNA. Nevertheless, recent reports showed direct involvement of the MOF Acetyl Transferase, a known chromatin structure modulator (Chatterjee et al. 2016) as well as members of the SIRTUIN family (Nakamura et al. 2008), in mtDNA transcription regulation. Additionally, the mtDNA is thought to fold into transcription-related loops, which are likely regulated (Martin et al. 2005; Uchida et al. 2017). These pieces of evidence prompted us to hypothesize that the human mtDNA may have a chromatin-like packaging with a role in gene expression regulation.

Here, by analyzing DNase-I-seq experiments from more than 320 different human samples (the RoadMap and ENCODE consortia) we found 29 DNase Genomics Footprinting (DGF) sites that were common to ∼90% of the samples. These sites exhibited a strong signature of negative selection, and were identified in mouse DNase-I-seq experiments, thus supporting their functional importance. Finally, nearly a third of the DGF sites precisely overlapped known mtDNA transcription regulatory elements and transcription pausing sites, thus strongly suggesting their involvement in the transcription machinery. We thus show first clues for a highly regulated organization of the mammalian mitochondrial genome.

## Results

### A common pattern of mtDNA DGF sites in a variety of human cell types

As a first step in characterizing mtDNA protein-DNA organization, we analyzed the comprehensive collection of DNase-I-seq experiments from the ENCODE consortium. We used only cells with experimental duplicates (70 cell samples), separately analyzed each of the duplicates, and retained only those DGF sites that were shared by the duplicates for further analyses. Briefly, to identify DGF sites we slightly modified a previously used approach (Mercer et al. 2011), and calculated an F-score for each mtDNA position in sliding sequencing read windows of variable size with a maximum of ∼124 bases, and a minimum of 18 bases (see below); each sliding window was divided into two flanking and one central fragments (i.e., a left ‘L’, right ‘R’ and central ‘C’, fragments-see details in Materials and Methods).

Our results revealed an average of 116.27 mtDNA DGF sites per cell line (SD=22.65), encompassing a mean of 1,868.36 bases (11.28%, +/-1SD=403 bases) of the mtDNA sequence. Although a total of 246 DGF sites were identified, covering more than half of the mtDNA sequence (8,660 bases, representing 52.27% of the mtDNA), more than half of the sites (N=135) were identified in ≥25% of the cell lines, suggesting that many mtDNA DGF sites are cell line-specific. While focusing on the most common DGF sites, we found that 61 were shared by more than 75% of the tested samples, of which 32 were identified in more than 90% of the samples (Fig. 1, Supplementary Table S1).

**Figure 1:**
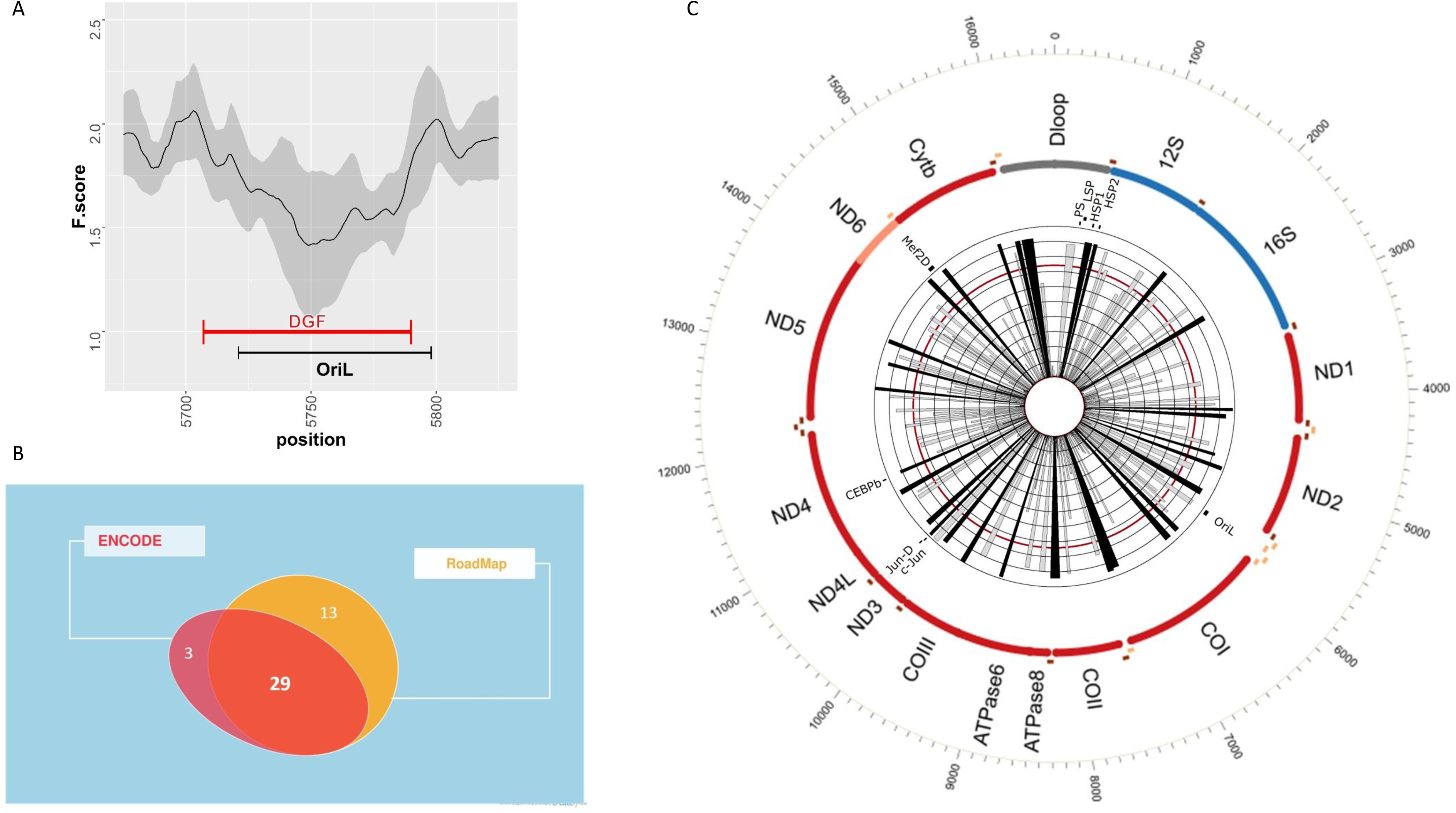
Histogram map of DGF patterns. A. Number of DGF sites identified in the ENCODE and RoadMap datasets, and their overlap. B. mtDNA location of all identified human DGF sites (n=246) according to their prevalence in the ENCODE sample collection. The inner ray histograms represent the prevalence of each site across ENCODE samples, with each circle marking 10^th^ percentile increments of the dataset (from 0-100%). The red circle marks 74% of the samples, which refers to the average + 1 STDV of the samples tested. Black rays indicate the most prevalent DGF sites.

To further validate the most abundant DGF sites, we extended our analysis to DNase-seq experiments generated by the RoadMap Consortium, comprising 264 human samples from various tissues, mostly fetal (n=224). In total, we identified 221 DGF sites in the RoadMap collection (Supplementary Table S1), of which 114 were in less than 25% of the samples and 64 were common to at least 75% of the tested samples. While focusing on the most abundant DGF sites, we found 42 sites that were common to more than 90% of the tested samples. Of the latter, 29 sites were located throughout the mtDNA shared by the ENCODE and Roadmap datasets, suggesting the presence of a number of highly conserved mtDNA DGF sites across adult and fetal human cell types.

As another validation of the identified DGF sites, we analyzed Assay for Transposase-Accessible Chromatin (ATAC-seq) experiments, which, in similar to DNase-seq, identify DNA sites that are occupied by proteins (Buenrostro et al. 2013). Our analysis of publically available ATAC-seq experimental data from three different human cell types (GM12878, neural stem cell, CD34+) clearly verified all of the 29 common DGF sites (not shown), thus supporting our approach and corroborating the true identification of the most common DGF sites in the human mtDNA.

### Nuclear mitochondrial DNA fragments (NUMTs) are not enriched in mtDNA DGF sites

One could argue that many of the mtDNA DNase-seq reads are contaminated by mtDNA fragments that were gradually transferred into the nuclear genome during the course of evolution, known as Nuclear Mitochondrial pseudogenes (NUMTs) (Hazkani-Covo et al. 2003; Mishmar et al. 2004). Notably, DGF sites are defined by reduced number of reads at a given site, which might be affected by excess of reads that were mapped to both the nucleus and to the active mtDNA. We therefore conducted a comprehensive screen to assess the amount of NUMT-associated reads in the ENCODE DNase-seq data, per sample per mtDNA position. To facilitate such a screen we used the collection of NUMT variants in the entire human genome (n=8031, see Material and Method), generated by Li and colleagues (Li et al. 2012). In general, the proportion of reads harboring NUMT variants comprised an average of only 0.165% of the reads (Fig. 2A, SD=0.668%). Furthermore, the proportion of NUMT reads was not statistically different between DGF and non-DGF sites across the entire human mtDNA (Fig. 2B, DGF sites=0.239%, SD=1.676%; non-DGF=0.163%, SD=0.304%). We conclude that NUMT reads had only negligible impact on our DGF analysis.

**Figure 2.**
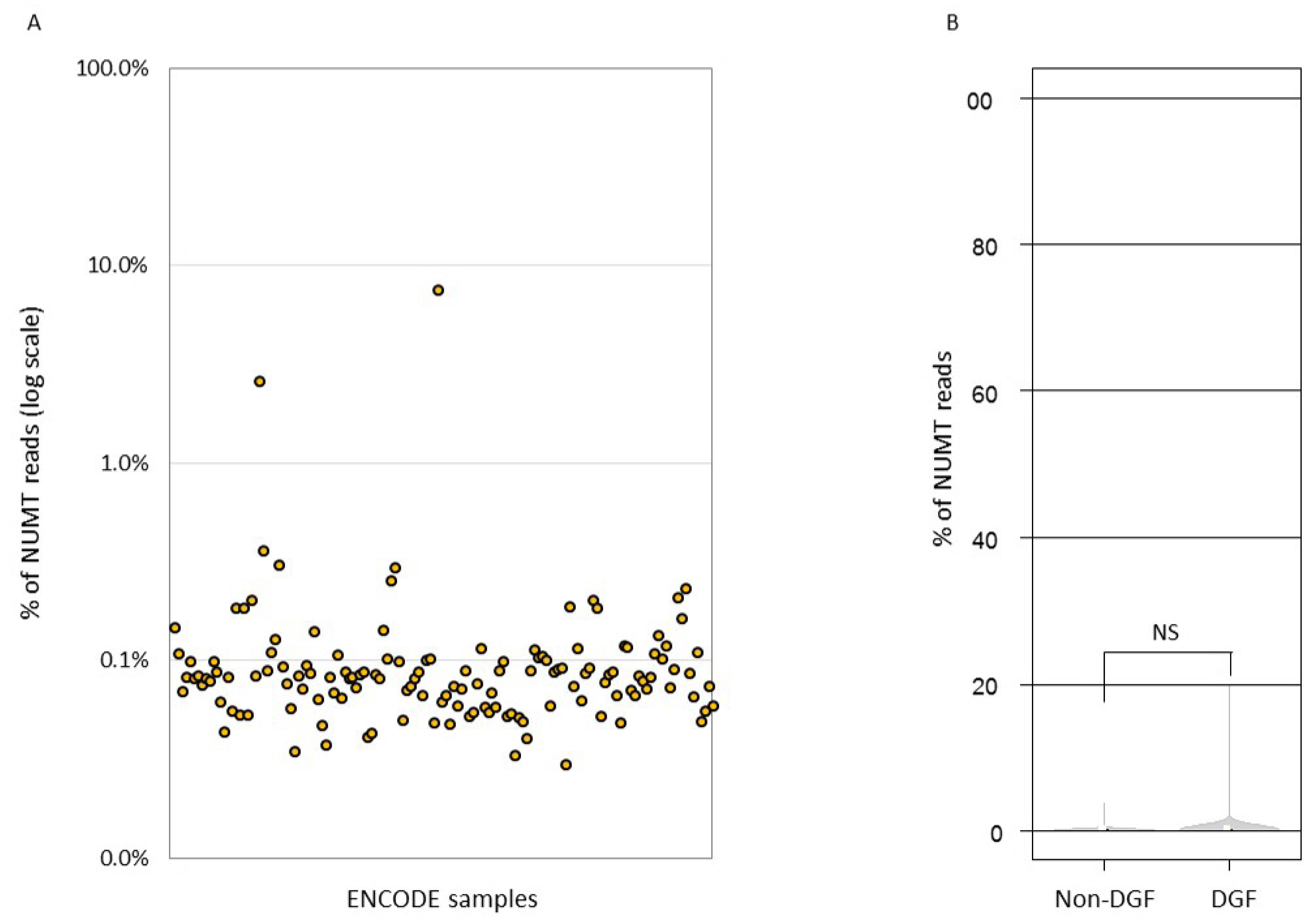
NUMT analysis. NUMT analysis was applied to all of the analyzed ENCODE samples. A. proportion of NUMT reads from the entire reads of each samples. Y-axes: Percentage of NUMTs reads from the total reads in each of the tested samples. X-axes: Each dot represents a single sample. B. A diagram comparing the prevalence of NUMT reads in DGF and non-DGF mtDNA sites.

### Tissue specific mtDNA DGF sites in the RoadMap collection

As many of the identified mtDNA DGF sites were shared only by subsets of the tested cell lines we hypothesized that at least part of those were tissue specific. To test for this hypothesis, we applied a PERMANOVA test and Principal Coordinates Analysis (PCO, using Primer-E -www.primer-e.com) to the RoadMap cell line dataset. Inspection of the entire dataset revealed strong clustering of samples from skin (see PCO in Fig. 3B, and p-values of the PERMANOVA test in supplementary Table S2). The same pattern was observed even while applying the same analysis to samples sharing the same developmental day (Fig. 3C). Nevertheless, we did not identify any additional cell type-specific clustering, suggesting that the observed variation in the prevalence of DGF sites could only partially be attributed to tissue specificity.

**Figure 3.**
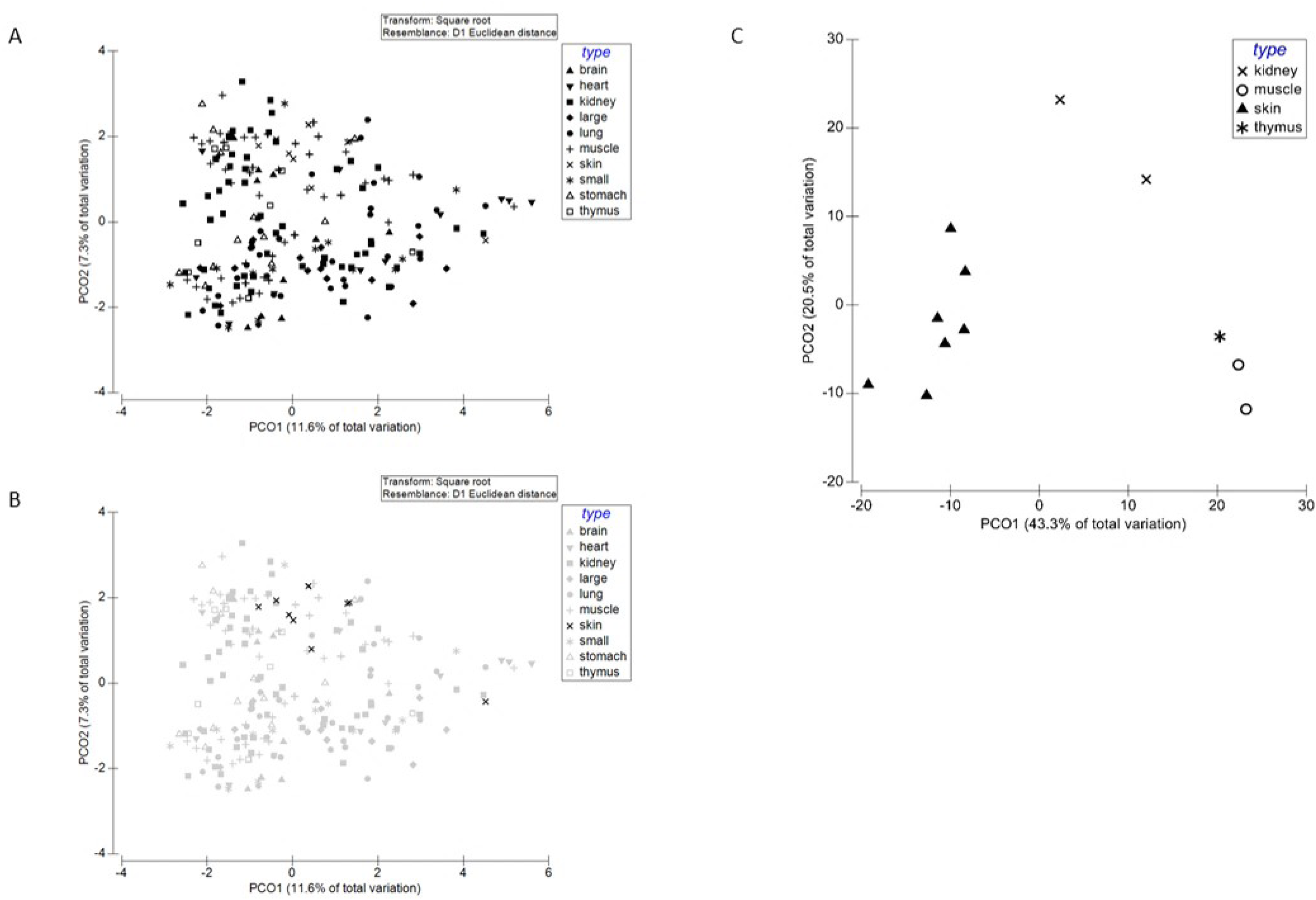
Evidence for tissue specificity in the mtDNA DGF pattern. PCO analysis of the RoadMap collection. A. All RoadMap samples. B. Skin samples. C. Samples from fetal day 97 (total=12; skin=7).

### The common mtDNA DGF sites co-localize with regulatory elements of mtDNA function

The existence of a common set of 29 mtDNA DGF sites shared between all available cell and tissue samples raise the possibility that sites are functionally important. As an initial step towards assessing such possibility, we screened for association between the common DGF sites and mtDNA elements with functional importance. This revealed that a subset of the DGF sites co-localized with heavy strand promoters 1 and 2, origin of replication of the light strand (OriL), the termination association site (TAS) and the recently identified protein-binding sites of c-Jun, Jun-D and CEBPb(Fig. 1). Although this finding supported the importance of DGF sites in the mtDNA, it raised a question regarding the functionality of the remaining DGF sites. Our recent identification of two transcription pausing sites that were common to 11 human cell lines tested (Blumberg et al. 2017), and their association with adjacent DGF sites, urged us to extend our screen to additional such sites throughout the mtDNA. This screen revealed 20 pausing sites (PS) that were shared by at least 8 of the 11 cell lines tested (Fig. 4, Supplementary Table S3), of which 17 sites were in the light strand and 3 in the heavy strand. Strikingly, 15 of these pausing sites either overlapped, or were in the immediate vicinity (less than 40bp) from DGF sites present in >75% of the cell lines. This suggests that many of the mtDNA DGF sites, and the factors that bind them, are involved in mtDNA transcription.

**Figure 4.**
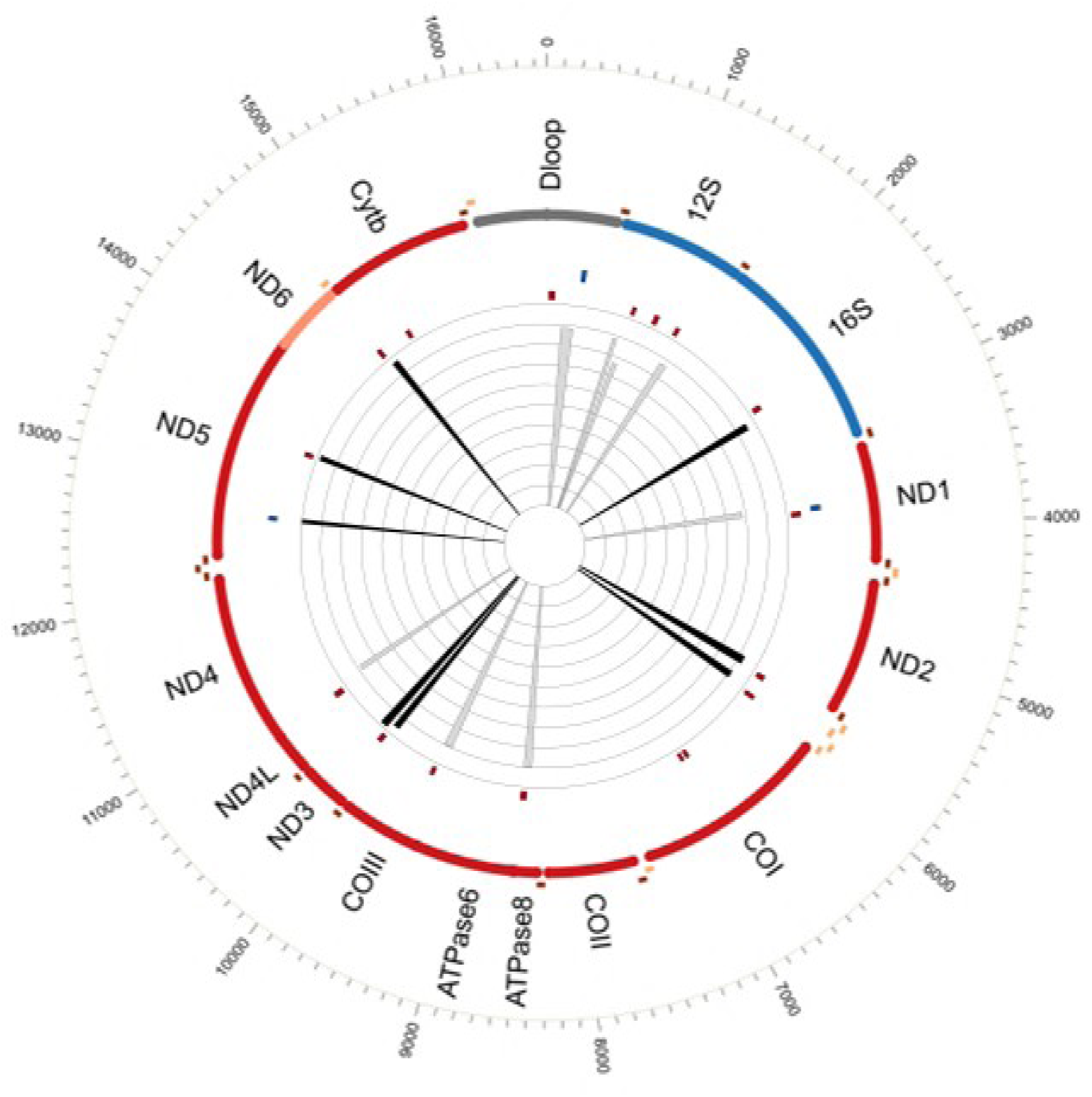
The common mtDNA DGF sites co-localize with regions of transcription pausing. The location of the 20 pausing sites across human mtDNA. Heavy strand (n=17) and light strand (n=3) pausing sites are marked by red and blue bars, respectively. The rays in the inner circle correspond to DGF sites present in >75% of samples, with the most common DGF sites colored in black.

### The mtDNA DGF pattern does not correlated with TFAM-binding sites

It is possible that most of the DGF sites result from increased affinity of the coating protein TFAM to certain mtDNA sites. To test for this possibility, we took advantage of available TFAM ChIP-seq experiments performed in HeLa cells (Wang et al. 2013). This analysis revealed 103 human mtDNA sites enriched for TFAM binding, as well as 88 sites depleted of TFAM ChIP-seq signals (marked as ‘high TFAM’ and ‘low TFAM’, respectively, in Fig. 5 and in Supplementary Table S4). Interestingly, we found significant enrichment of the HeLa cells DGFs (including the 29 common DGF sites) at the ‘low TFAM’ sites (36 out of 88 sites, p = 0.0099, one tail Fisher's exact test), as opposed to the ‘high TFAM’ sites (16 out of 103, p = 0. 8435, one tail Fisher's exact test). These results indicate that most DGF sites are unlikely occupied by TFAM (Fig. 5).

**Figure 5:**
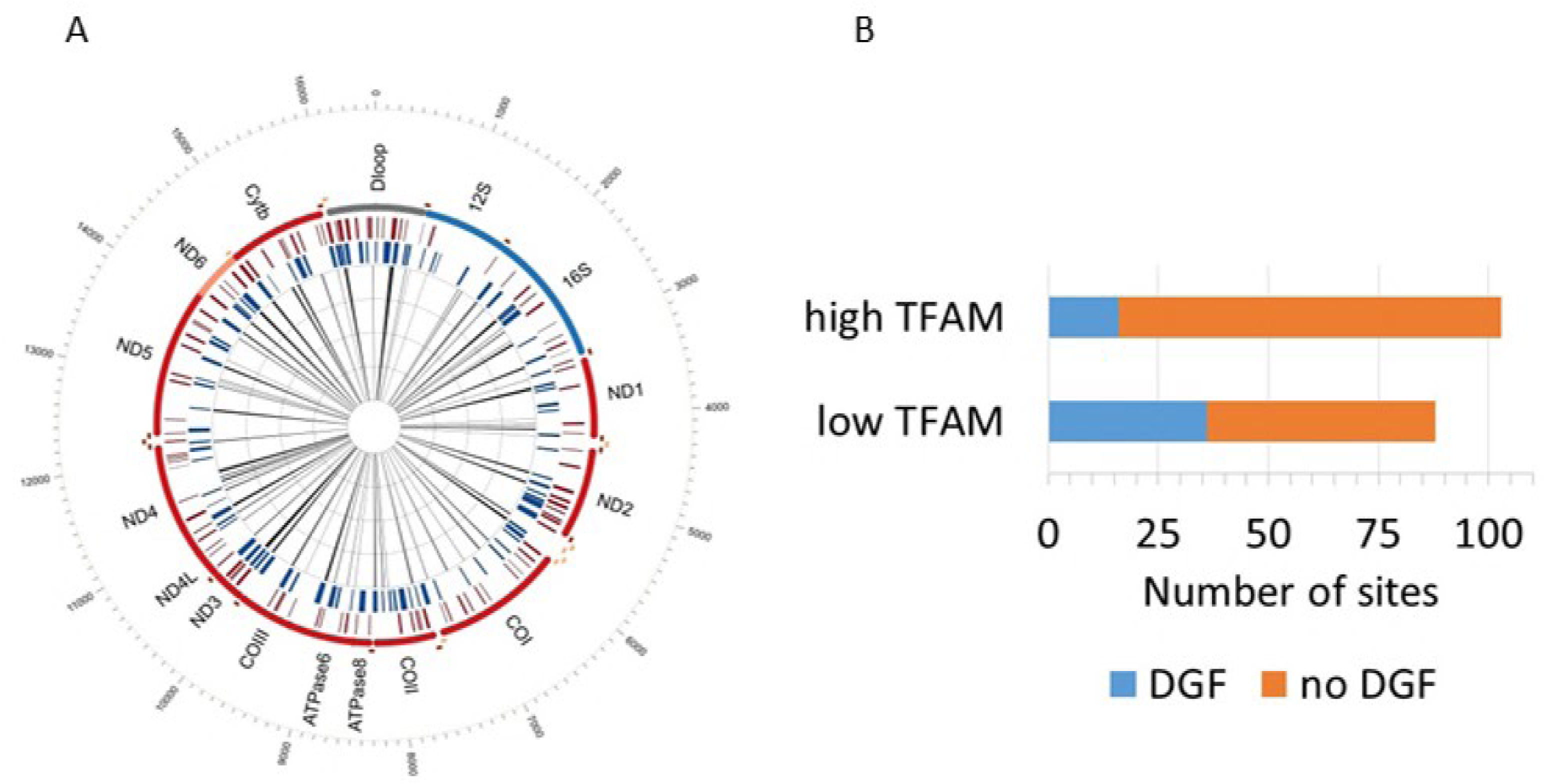
TFAM binding and mtDNA DGF pattern in HeLa cells. A. An mtDNA map of the DGF sites in HeLa cells (inner histograms), along with the low-TFAM (blue bars) and high-TFAM sites (red bars). B. The proportion of high and low TFAM sites among mtDNA DGF sites in HeLa cells. Y-axis – the group of tested TFAM sites. X-axis – number of sites. DGF sites are colored in blue and non-DGF sites are in red.

### Negative selection and evolutionary conservation of the common mtDNA DGF sites

The high frequency of genetic variants throughout the human mtDNA, and the knowledge of their prevalence in human worldwide populations enabled using SNP density as a tool to identify signatures of selection (Levin et al. 2013). Although DGF sites were identified throughout the mtDNA, including protein-coding sequences, we limited the assessment of SNP density to 3^rd^ codon positions (n=428) in the 23 DGF sites within protein-coding genes; this focus was aimed to avoid misinterpretation due to selection acting on protein-coding sequences. Notably, DGFs within RNA genes (tRNAs and rRNAs) were not included in this analysis as there was not an easy way to distinguish the selective constraints acting on the gene sequences from that acting on the DGFs. Our calculated number of ancient variants in protein-coding DGF sites (3^rd^ codon positions only, 23 DGF sites) was significantly lower than the entire set of 3^rd^ codon positions in protein-encoding gene sequences throughout the mtDNA (p=0.0486, tested in 1,000 simulation replicates, see Materials and Methods), consistent with purifying selection acting to constrain the mtDNA sequence in DGF sites. To gain further insight into the signature of selection in mtDNA DGF sites, we estimated the ratio between ancient mutations (nodal mutations, which were already exposed to natural selection for a long time) and relatively recent mutations (present at the phylogenetic tree tips, and had less time to undergo selection) as previously performed (Blumberg et al. 2014). We found a significantly lower ratio in DGF sites as compared to control (p = 0.0066, tested in simulation, see Materials and Methods). This finding suggests that DGF sites are less likely to sustain older mutations, thus further implying that mtDNA DGF sites are negatively selected.

Since evolutionary conservation reflects functional importance, we assessed the conservation of DGF sites from man to mouse by analyzing mtDNA DGF sites in mouse DNase-seq data generated by the ENCODE consortium from 43 cell types in experimental duplicates (Supplementary Table S5). We found that 89.66% of the most common human mtDNA DGF sites were in the same mtDNA regions harboring mouse DGF sites in at least 10% of the mouse cell lines tested (Fig. 6). Taken together, these data support the functional importance of mtDNA DGF sites during evolution.

**Figure 6:**
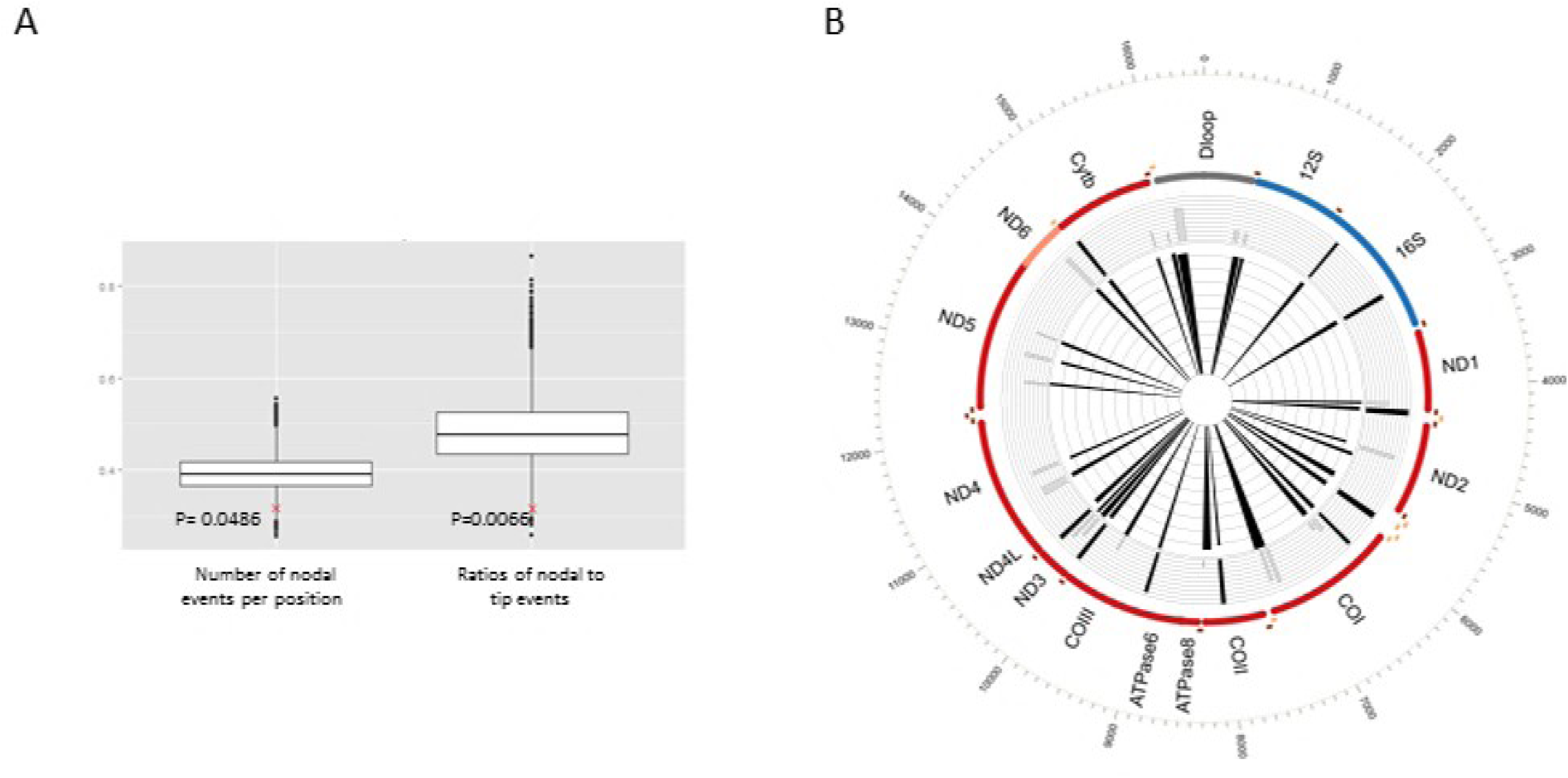
Conservation of common mtDNA DGF sites. A. Boxplot graphs of the SNP density simulation. Y-axis - arbitrary units. X-axis – SNP density tests. The left box plot – number of nodal events per position. The right box plot – ratio of nodal to tip mutational events at the DGF sites tested. P-values are indicated. B. Common human DGF sites across the human mtDNA map are presented, while super-imposing mouse mtDNA DGF sites. The inner ray histogram corresponds to the 29 most common human mtDNA DGF sites, while the outer histogram corresponds to the mouse mtDNA DGF sites in grey-scale, with black representing the most common sites.

To get a first glance into the physiological importance of mtDNA DGF sites, we screened through ENCODE DNase-seq data from human cells exposed to diverse physiological conditions to identify conditions that were previously found to affect mitochondrial function. Treatment of mammalian cells by interleukin-3 or erythropoietin was previously shown to impact mitochondrial activity (Carraway et al. 2010; Wellen et al. 2010). Our results indicate that treatment of human hematopoietic CD34+ lymphocytes by interleukin-3, hydrocortisone, succinate and erythropoietin led to a 100 nucleotides shift of the DGF site at mtDNA positions 365-398 to nucleotide positions 279-347 (Fig. 7). As this treatment led to gain of a new mtDNA DGF site within Conserved Sequence Block II (CSBII), a known mtDNA transcription-to-replication transition point (Pham et al. 2006), we calculated mtDNA copy numbers in the treated and control cells. This analysis revealed a two-fold increase in mtDNA copy numbers in the treated cells, suggesting a functional impact for the DGF site shift (Fig. 7). We noticed that the experiments performed by ENCODE used control and treated cells having different mtDNA genetic backgrounds (haplogroups): whereas the control cells belonged to either the U5b1 or K1a2a haplogroup the treated cells belonged to the T2e haplogroup. To control for the possible inherent differences in mtDNA copy number between the mentioned mtDNA haplogroups, we re-analyzed the mtDNA copy number of B-lymphocytes from phylogenetically related genetic backgrounds, extracted from the 1000 Genomes Project (Cohen et al. 2016). Our results indicate no significant difference between the mtDNA copy number of cells belonging to mtDNA haplogroup U5 (N=61 samples, mean mtDNA copy number=421.77, SD= 88.36), haplogroup K1 (n=15 samples, mean mtDNA copy number=416.46, SD=85.58) and haplogroup T2 (n=18 samples, mean mtDNA copy number=441.8, SD=112.79 (T-test; U5 vs K1 - p=0.837 (NS); U5 vs T2 - p=0.695 (NS); T2 vs K1– p =0.636 (NS)). Taken together, our analyses offer first clues for physiological response of our discovered mtDNA DGF sites.

**Figure 7:**
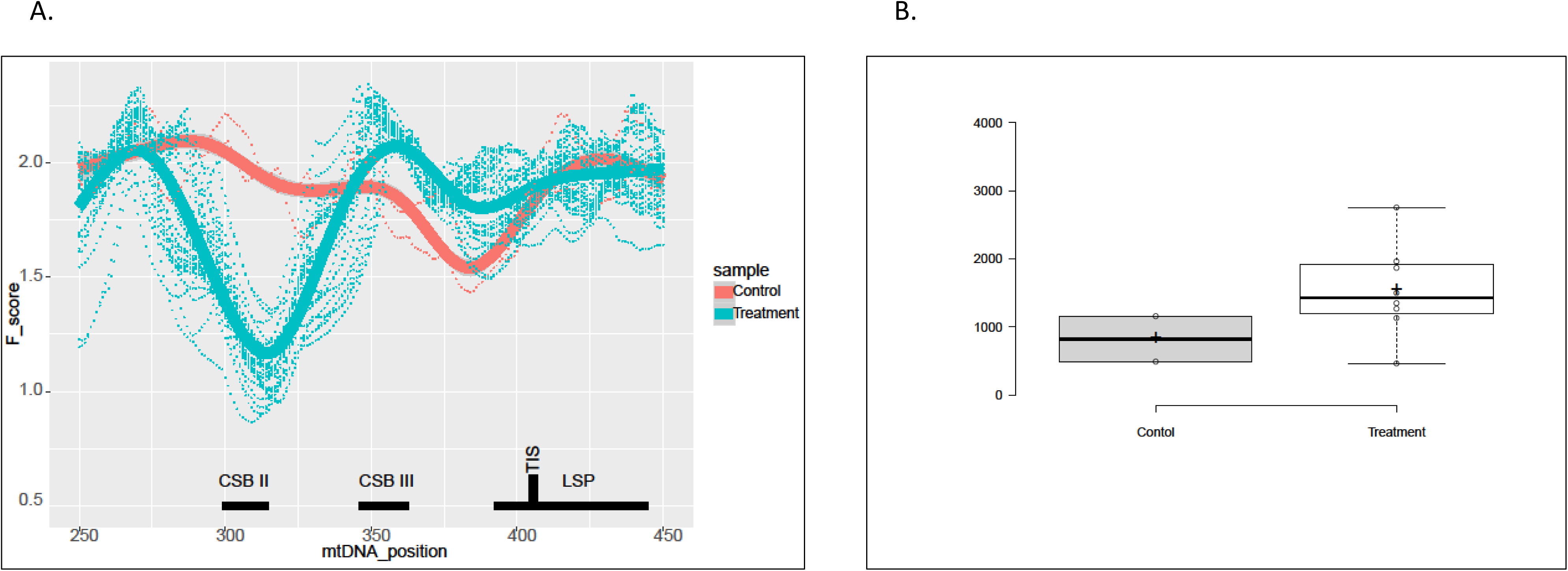
Gain of mtDNA DGF site at the transcription-replication switch site, upon CD34+ IL-3 dependent activation. A. Representation of the F-scores around the Light strand transcription start site. Y axes: F-score, X-axes: mtDNA position. Lowest F-score – the most DNase I protected region. Thick line – average of F-scores across the analyzed samples. Dotted line – F-score calculated for single experiments. Red – control experiment, gray – experiments of treated cells. B. mtDNA copy number in control and treated cells.

## Discussion

In the current study, we demonstrate for the first time, that the vast majority of human cell types display a conserved mtDNA footprinting pattern. Specifically, our analysis of DNase-I-seq experiments in more than 320 human cell types revealed 29 DGF sites which were common to >90% of the tested samples. Since mtDNA DGF sites coincided with lower TFAM occupancy, higher-order organization of the human mitochondrial genome likely involves proteins other than TFAM. This observation is consistent with the argument that human mtDNA is not evenly coated by TFAM (Rebelo et al. 2009). Therefore, although mtDNA condensation increases with elevated cellular concentration of TFAM (Kukat et al. 2015), mtDNA packaging is likely more organized and more complex than once thought. The striking co-localization of the mtDNA DGF sites with transcription pausing sites and regulatory elements of mtDNA gene expression, along with the strong negative selection that govern these DGF sites, suggest that mitochondrial DGF sites have functional importance, most probably for mtDNA transcription.

Our finding that the mtDNA DGF sites are likely under-occupied by TFAM suggests a role for another protein/s, yet to be characterized. Previously, several proteins have been assigned to the mitochondrial nucleoid, yet only TFAM was clearly shown to participate in mtDNA packaging (Lee and Han 2017). Our highly ordered DGF site pattern in human cells implies departure from current thought – it suggests a ‘chromatin’-like structure in the human mtDNA. Our observed correlation between the mtDNA DGF sites and transcription regulatory sites tempts us to suggest a conceptual similarity between the higher order organization of the nuclear and mitochondrial genomes, hence reflecting a regulatory aspect of adaptation of the mitochondrion to its ancient host. The growing collection of transcription factors (and other DNA-binding proteins) that are used in ChIP-Seq experiments increase the odds of identifying this unknown mtDNA-binding factor/s in the future.

While considering both rare and common mtDNA DGF sites, profound variability in their prevalence and pattern among cell types was observed. In fact, most mtDNA DGF sites were identified in less than 25% of the tested samples (135/245 DGF sites). Although a skin-specific pattern was observed, most variability of mitochondrial DGF site patterns could not be explained by tissue specificity, or fetal versus adult tissue differences (not shown). One possibility to explain such variability stems from differences in physiological conditions. Indeed, by targeting bacterial DNA methyltrasferases to the mitochondria in human cells it was found that the level of mtDNA protein occupancy differed among mtDNA regulatory sites in response to various physiological insults (Rebelo et al. 2009). It is thus plausible that sample variability in patterns of mtDNA DGF sites could in part reflect differential responses to physiological differences. This is supported by our observed shift in the DGF site pattern in CD34+ cells treated with interleukin-3, hydrocortisone, succinate and erythropoietin as compared to control (Fig. 7). This strongly suggests that our observed DGF pattern cannot be attributed to recently suggested sequence-specific DNase I cutting bias (He et al. 2014), but rather has biological importance.

In summary, our comprehensive analysis of human mtDNA DNase Genomic Footprinting experiments revealed 29 common negatively selected mtDNA DGF sites that were common to 90% of ∼320 different cell types. This reflects a systematic and regulated organization of this genome in the vast majority of human tissues. The over-representation of such DGF sites in mtDNA regions with lower TFAM occupancy suggests that mtDNA-binding factors other than TFAM are involved in this mtDNA-protein organization. Secondly, we found tissue-specific patterns of mtDNA DGFs, suggesting their physiological importance. Thirdly, most transcription pausing sites throughout the mitochondrial genome co-localized with mtDNA DGF sites, thus supporting the involvement of the latter in progression of mtDNA transcription. Hence, packaging and regulation of mitochondrial genome transcription are far more regulated, and certainly more complex, than once thought.

## Materials and Methods

### Sample-specific mtDNA sequence re-construction and mapping; Coverage calculation; Circular-like mapping of sample-specific mtDNA sequence

Analyses were applied as described previously (Blumberg et al. 2017). In brief, after mapping the sequencing reads to the mtDNA reference genome (rCRS - NCBI Reference Sequence: NC_012920.1), sample-specific mtDNA reference genome was re-constructed and the DNase-seq reads were aligned while taking into account the circular organization of the mtDNA as recently performed (Blumberg et al. 2017).Read coverage for each position was calculated using the 'genomecov' command in BEDtools (http://bedtools.readthedocs.org/en/latest/ version 2.25) (Quinlan and Hall 2010).

### DNase-seq analysis

ENCODE DNase-seq fastq files were downloaded from the ENCODE consortium (The-ENCODE-Project-Consortium 2012) website (http://hgdownload.cse.ucsc.edu/goldenPath/hg19/encodeDCC/wgEncodeUwDnase/) for human and http://hgdownload.cse.ucsc.edu/goldenPath/mm9/encodeDCC/wgEncodeUwDnase/ for mouse. ROADMAP DNase-seq SRA files were downloaded from the ROADMAP epigenomics project website (Roadmap Epigenomics et al. 2015) (http://egg2.wustl.edu/roadmap/web_portal/index.html). Data of treated and control CD34+ cells was taken ENCODE (http://www.encodeproject.org/search/?searchTerm=hematopoietic%2Bmultipotent%2B www.encodeproject.org/search/?searchTerm=hematopoietic+multipotent+progenitor+cell&type=Experiment&assay_title=DNase-seq). DGF sites were identified following the method outlined in: github.com/StamLab/footprinting2012/.Briefly, for each mtDNA nucleotide position, an *F*-score was calculated in sliding read windows of variable size with a maximum of 124 bases, and a minimum of 18 bases (see below) using the following equation: F = (C + 1)/L + (C + 1)/R, where ‘C’ represents the average number of reads in the central fragment C, ‘L’ represents the average read count in the proximal fragment, and ‘R’ represents the average read count in the distal fragment. Following published specifications (github.com/StamLab/footprinting2012/) (Neph et al. 2012), the variable sliding window sizes was comprised of 6-100 bases for the ‘C’ fragment, and 6-12 bases for each of the ‘L’ and ‘R’ fragments. Regions showing the lowest F scores, which reflected relative depletion of reads, were considered DGF sites.

### Pausing sites analysis

Pausing sites were identified as we recently described (Blumberg et al. 2017). In brief, Pausing Index (PI) was calculated using the following equation: PI = (T+1)/(GB+1), where T represents density of reads in 20 bases of the tested position and GB represents the density of reads in the gene body. To minimize putative reciprocal influence of close internal pausing sites, 'gene body' of each position was calculated in sliding windows of 10-1,000 bases that flank each of the tested positions (both upstream and downstream), and the highest value for each position was considered as the optimal value for the tested position. For each experiment, positions exhibiting higher PI values than the average plus 1 SD were considered as pausing sites.

### NUMTs analysis

Proportion of NUMT reads was estimated by counting mtDNA mapped DNAse-seq reads (within BAM files) that contain NUMT variants using bam-readcount (https://github.com/genome/bam-readcount). For the sake of consistency, without compromising the comprehensive nature of our analysis we used an updated collection of 8031 human NUMT variants published by Li et al (Li et al. 2012). For every analyzed sample, sample-specific NUMT collection was generated by screening the reconstructed sample-specific mtDNA sequences.

### Assessment of SNP frequency at the mtDNA DGF sites

Phylogenetic analysis of nearly 10,000 whole mtDNA sequences representing all major global populations allowed for the extraction of multiple ancient mutational events (Levin et al. 2013). The number of mutational events at the 3^rd^ codon position of coding region DGF sites was counted and compared to the number of mutational events in the entire set of 3^rd^ codon positions in all mtDNA protein-coding genes. The logic underlying this analysis was based on the following simple argument: A reduced number of mutational events at a given tested site reflects a signature of negative selection. To confirm that the lower number of variants in the 3^rd^ codon position in the DGF sites cannot be explained by chance, we applied 10,000 replicated simulations in which the sample size of the 3^rd^ codon position within mtDNA DGFs was retained but shuffled with sites throughout the mtDNA protein-coding region.

### Identification of mtDNA sites with high and low TFAM occupancy

We re-analyzed the results of four biological replicates of TFAM ChIP-seq experiments in HeLa cells (Wang et al. 2013). Although TFAM binds throughout the mtDNA, we identified TFAM enriched and low occupation sites by calculating F-scores: For TFAM enriched sites, the top F-scores were taken in account, whereas the lowest F-scores were considered sites with low TFAM occupancy (similar to the identification of DGF sites).

### PCO and PERMANOVA analyses

PCO and PERMANOVA analyses were performed to assess possible tissue specific pattern of DGF sites. These analyses were applied using Primer-E (version 6 http://www.primer-e.com/NUMTs) while applying default parameters.

### Visual representations of the mitochondrial genome

CIRCOS was used for visualization of all circular mtDNA graphs (Krzywinski et al. 2009).

### MtDNA copy number estimation

MtDNA copy number of DNase-seq experiments was estimated as recently described (Cohen et al. 2016) with minor modifications. In brief, to control for over/under representation of sequencing reads in certain genomic regions we compared the mtDNA read numbers to regions of 5 x 10^6^ bases from each of the 22 autosomal chromosomes.

### Haplogroups assignment

mtDNA haplogroup assignment of samples was performed by analyzing reconstructed whole mtDNA sequences using Haplogrep (Kloss-Brandstatter et al. 2011).

## Figure Legends

**Supplementary Table S1: Human mtDNA DGF sites from ENCODE and RoadMap data.**

**Supplementary Table S2: RoadMap clustering statistical data.**

**Supplementary Table S3: Common Pausing sites throughout the Human mtDNA.**

**Supplementary Table S4: TFAM occupancy compared to DGF pattern in HeLa cells**.

High and low TFAM occupancy, as a result of TFAM ChIP-seq analysis performed in human HeLa cells. The specific mtDNA DGF pattern in HeLa cells is indicated.

**Supplementary Table S5: Mouse mtDNA DGF sites extracted from the ENCODE data.**

